# Both CD8 and CD4 T cells contribute to immunosurveillance preventing the development of neoantigen-expressing autochthonous sarcomas

**DOI:** 10.1101/2023.04.04.535550

**Authors:** Jonathon E. Himes, Amy J. Wisdom, Laura Wang, Sam J. Shepard, Andrea R. Daniel, Nerissa Williams, Lixia Luo, Yan Ma, Yvonne M. Mowery, David G. Kirsch

## Abstract

The adaptive immune system plays an essential anti-tumor role through immunosurveillance and response to immunotherapies. Characterizing phenotypic features and mechanisms of dysfunction of tumor-specific T cell populations may uncover novel immunotherapeutic targets and biomarkers of response. To study tumor-specific T cell responses *in vivo*, a tumor model must express a known neoantigen. While transplant models with known neoantigen expression are widely available, autochthonous tumor models in which the tumor coevolves with the immune system are limited. In this study, we combined CRISPR/Cas9 and sleeping beauty transposase technology to develop an autochthonous orthotopic murine sarcoma model with oncogenic Kras^G12D^, functionally impaired p53, and expression of known MHCI and MHCII sarcoma neoantigens. Using MHC tetramer flow cytometry, we identified a tumor-specific immune response in the peripheral blood as early as 10 days after tumor induction leading to tumor clearance. Tumors developed at high penetrance after co-depletion of CD8 and CD4 T cells, but depletion of either CD8 or CD4 T cells alone was insufficient to permit tumor growth. These results suggest that CD8 and CD4 T cells can independently contribute to immunosurveillance leading to clearance of sarcomas expressing MHCI and MHCII neoantigens.

## Introduction

Even though the immune system can prevent cancer through immunosurveillance, roughly one-third of Americans will develop cancer in their lifetimes^1^. Immune evasion by tumor cells can occur via tumor-intrinsic or immune cell-specific mechanisms^2^. Tumor cells can be directly killed by cytotoxic CD8 T cells. Therefore, cytotoxic CD8 T cells are widely accepted as important contributors to the anti-tumor function of the immune system. Cytotoxic T cells are a target for many of the most successful immunotherapies to date, such as immune checkpoint inhibitors^3^. However, the importance of CD4 T cells for anti-tumor responses is less clear with conflicting reports in the literature^4–8^. This uncertainty regarding the potential anti-tumor role of CD4 T cells stems from the diverse functionalities of different CD4 T cell subsets, such as helper T cells and immunosuppressive regulatory T cells. For example, several studies have demonstrated a pro-tumor effect mediated by CD4 T cells^4–6^. Regulatory CD4 T cells in tumor-draining lymph nodes can promote immune tolerance of the tumor^6^. In contrast, other studies have reported that CD4 T cells can aid in cytotoxic CD8 T cell activation as helper cells^9^. In several transplant tumor models in mice, the anti-tumor potential of CD4 T cells has been clearly demonstrated^7,10^. Adoptive transfer of CD4 T cells has been shown to elicit impressive anti-tumor responses against several transplanted tumor models, even rivaling the effectiveness of CD8 T cell adoptive transfer^7^. Similarly, a study demonstrated that when sarcoma cell lines expressing either a known MHCI-presented antigen or an MHCII-presented antigen were injected into the flank of syngeneic mice, tumors grew and did not respond to anti-PD-1 therapy^10^. However, when sarcoma cells expressing both an MHCI-presented antigen and an MHCII-presented antigen were injected into syngeneic mice, they elicited a CD8 and CD4 T cell response which limited tumor penetrance *in vivo*, and the tumors that formed were responsive to anti-PD-1 therapy^10^.

While these data support the concept that CD4 T cells play a major anti-tumor role complementary to CD8 T cells, these studies are limited by the model systems utilized. Preclinical trials utilizing transplant tumor models have demonstrated promising therapeutic responses to treatment modalities such as the combination of immune checkpoint blockade and radiation therapy^11–14^, but these responses have not subsequently translated into similar outcomes in clinical trials^12,15,16^. This disparity may be due to differences in the tumor microenvironment given that transplant models lack the gradual tumor-immune system coevolution that occurs in murine autochthonous tumors and in human cancers. Therefore, we developed a novel autochthonous sarcoma model presenting previously defined MHCI and MHCII tumor neoantigens^10,13^ to the immune system in order to evaluate the impact of CD8 T cells and CD4 T cells in autochthonous tumor development and in immune-mediated autochthonous tumor clearance.

## Methods

### Mice

All animal studies were performed in accordance with protocols approved by the Duke University Institutional Animal Care and Use Committee (IACUC) and adhere to the NIH Guide for the Care and Use of Laboratory Animals^17^. All mice used in the study were C57BL/6J wildtype mice obtained from Jackson Laboratory C57BL/6J (JAX #000664). Groups of mice for each experiment were age- and sex-matched to minimize effects of sex- or age-mediated differences.

### Sarcoma initiation

Autochthonous sarcomas were induced in 6-12 week old C57BL/6J mice using *in vivo* electroporation (IVE) of a two plasmid system. IVE of plasmids into muscle cells of the gastrocnemius was conducted as previously described^18^. Briefly, 50ug of each plasmid diluted to 2mg/mL in normal saline were injected into the gastrocnemius with an insulin syringe. Then needle electrodes were used to deliver six total 200ms pulses of 100V directly to the muscle (Electro Square Porator ECM830; BTX, San Diego, CA). All autochthonous sarcoma models described here utilized the same transient plasmid, which expressed the sleeping beauty transposase SB100x, a sgRNA targeting *Trp53* (sgp53 sequence: GATGGTAAGGATAGGTCGG), and Cas9 endonuclease. The second plasmid contained two transposase insertion sites to allow for insertion of a gene into the genome of cells. This integrating plasmid varied depending on the autochthonous model and contained oncogenic *Kras*^*G12D*^ alone (SBKP), mLama4 preceding a 2A sequence leading into *Kras*^*G12D*^ (mLama4 SBKP), or a carrier GFP protein containing the predicted core neoantigenic peptides of mLama4 (VGFNFRTL) and mItgb1 (VNGYNEAIVHVVETP) preceding a 2A sequence leading into *Kras*^*G12D*^ (mLama4/mItgb1 SBKP). Induced legs were measured with calipers 3 times weekly to monitor for tumor development. Mice were euthanized with CO2 if tumors reached more than 13mm in any dimension, per the Duke University IACUC guidelines.

Transplant sarcomas were induced in 6-12 week old C57BL/6J mice by injecting 100,000 cells in 50uL RMPI directly into the gastrocnemius muscle. Transplantation cell lines were derived from autochthonous tumors harvested at >150mm^3^. Harvested tumors were mechanically dissociated with the gentleMACS dissociator (Miltenyi Biotec) and chemically dissociated with the Miltenyi Tumor Dissociation Kit (Miltenyi Biotec) at 37C for 45 minutes. Dissociated cells were strained through a 70um filter and plated at 37C. Cell lines were passaged once prior to transplantation.

### Flow cytometry

Blood was harvested from animals by submandibular bleed with a 5mm lancet. Peripheral blood mononuclear cells (PBMC) were isolated using gradient centrifugation with Ficoll-Paque Plus (Cytiva). Blood diluted 2:1 in RPMI was overlayed onto Ficoll-Paque and centrifuged for 15 minutes at 350g. The buffy coat was aspirated and washed with RPMI. PBMCs were stained with a panel of fluorochrome-conjugated antibodies reactive with the following cell surface markers: CD45-APC/Cy7 (clone 30-F11; BD Biosciences), CD3-PE/Cy7 (clone 500A2; Biolegend, San Diego, CA), CD8-FITC (clone KT15; ThermoFisher Scientific, Waltham, MA), CD4-BV421 (clone GK1.5; BD Biosciences). Antibody titrations are available in **Supplemental Table 1**. PBMCs were also stained with Zombie Aqua (1:320 dilution; Biolegend, San Diego, CA) to identify dead cells. To identify T cells specific for mLama4 or mItgb1, PBMCs were stained with H-2K(b)-mLama4 (VGFNFRTL) or I-A(b)-mItgb1 (VNGYNEAIVHVVETP) MHC tetramers conjugated to PE and APC obtained from the NIH Tetramer Core Facility. Stained cells were acquired on a BD LSRFortessa Cell Analyzer at the Duke Human Vaccine Institute Flow Cytometry Core Facility. Compensation was accounted for using AbC Total Antibody Compensation Beads (ThermoFisher Scientific, Waltham, MA). Data were analyzed using FlowJo 10.

### Immune cell depletion

All T cell depletions were conducted with a single injection 3 days prior to using IVE for tumor induction. Anti-CD4 (InVivoMAb clone GK1.5; BioXcell, Lebanon, NH), anti-CD8 (InVivoMAb clone 2.43; BioXcell, Lebanon, NH), and/or isotype control (InVivoMAb rat IgG2b clone LTF-2; BioXcell, Lebanon, NH) antibodies were injected intraperitoneally at a concentration of 1mg/mL in a volume of 200uL in pH 7.0 InVivoPure buffer (BioXcell, Lebanon, NH).

### Soft agar transformation assay

Wildtype mouse embryonic fibroblasts (MEFs) were harvested from C57BL/6J mice and underwent *in vitro* electroporation of SBKP, mLama4 SKBP, or mLama4/mItgb1 SBKP plasmids using the Lonza 4D-Nucleofector per the manufacturer’s instructions. Soft agar assays were used to evaluate cell line transformation potential as previously described^19^. Briefly, 15,000 cells were plated in 0.4% bactoagar RPMI with 10% FBS and penicillin/streptomycin over a 0.6% bactoagar RPMI layer in 6 cm plates. Plates were incubated for 3 weeks to allow colony formation prior to staining with 0.05% crystal violet in 10% ethanol for 1 hour. Colonies were imaged using a Leica inverted light microscope and colonies were manually counted by an investigator blinded to treatment group.

### Statistical Analysis

Statistical tests performed for all experiments are indicated in the figure legends. Non-parametric T tests or ANOVA were employed to compare cell proportions between groups. Kaplan-Meier analysis with log-rank test was used to compare tumor penetrance between groups. Multiple comparisons were accounted for using the false discovery rate (FDR) corrections. All statistical analysis was conducted with Prism 9 (GraphPad).

## Results

### Generating autochthonous sarcomas with multi-plasmid in vivo electroporation

To generate an autochthonous sarcoma model with known neoantigens, we first used CRISPR/Cas9 and sleeping beauty transposase technology to develop a non-antigenic autochthonous sarcoma model where the plasmid backbone could be modified to add known neoantigens. Our previous work demonstrated that deleting both alleles of *p53* and activating oncogenic Kras^G12D^ in the muscle is sufficient to initiate the development of sarcomas with few nonsynonymous mutations and thus low neoantigenic load^20^. We developed a multi-plasmid IVE system capable of inducing non-antigenic sarcomas in C57BL/6J wildtype mice (**Figure 1A**). In this system, two plasmids are injected into the gastrocnemius muscle of mice prior to electroporation. One plasmid transiently expresses Cas9, a sgRNA targeting *Trp53*, and a sleeping beauty transposase (SB100x) capable of inserting large sections of DNA flanked by transposase insertion sites into the eukaryotic genome. Cas9 and sgRNA targeting *Trp53* induce insertions/deletions (indels) in both *Trp53* tumor suppressor alleles, rendering them non-functional. The second plasmid carries a single transcript encoding Kras^G12D^ that is flanked by transposase insertion sites so that *Kras*^*G12D*^ is inserted into the genome by the sleeping beauty transposase. This autochthonous sarcoma model (SBKP) was 100% penetrant with detectable tumors developing as early as 29 days after IVE (**Figure 1B**).

**Figure 1.**
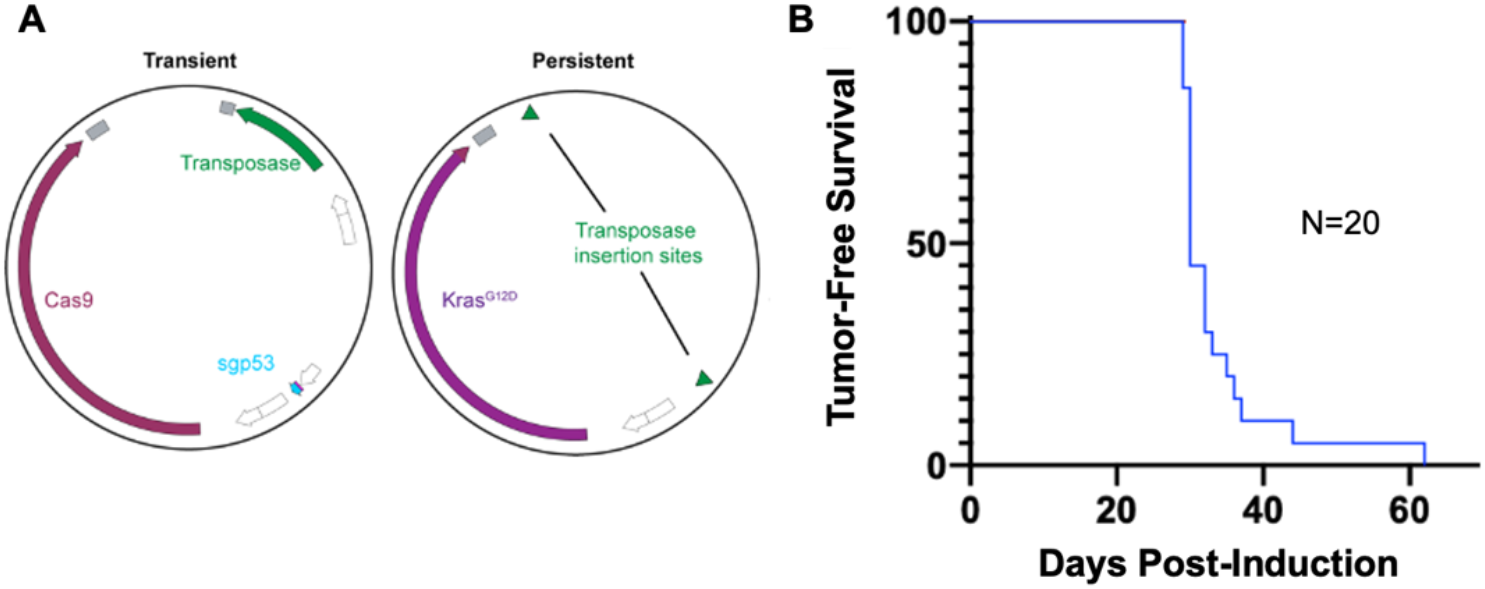
IVE of the SBKP plasmid system into the gastrocnemius of mice results in highly penetrant autochthonous SBKP sarcomas. **A)** Illustrations of the transient and persistent plasmids of the SBKP system injected into the gastrocnemius muscle prior to electroporation. White segments indicate promoter regions and grey segments indicate poly-A tails. The transient plasmid contains the sleeping beauty transposase SB100x, Cas9, and sgRNA targeting *Trp53* (sgp53). The persistent plasmid contains transposase insertion sites flanking Kras^G12D^. **B)** Kaplan-Meier curve of tumor-free survival after IVE of SBKP plasmids into the gastrocnemius of C57BL/6J wildtype mice.

### Engineering MHCI and MHCII neoantigen expression into autochthonous sarcomas

A major challenge to model neoantigen expression in autochthonous tumor models is immune clearance of highly immunogenic tumor-initiating cells, resulting in poor tumor penetrance. Therefore, we modified the integrating *Kras*^*G12D*^ DNA to include two neoantigens that had previously been identified in a 3-methylcholanthrene (MCA)-induced sarcoma cell line (**Figure 2A**)^10,13^. These two tumor neoantigens, *Lama4*^*G1254V*^ (*mLama4*) and *Itgb1*^*N710Y*^ (*mItgb1*), had been predicted with a bioinformatics algorithm and validated experimentally in this MCA-driven sarcoma cell line as MHCI-presenting and MHCII-presenting neoantigens, respectively^10,13^.

**Figure 2.**
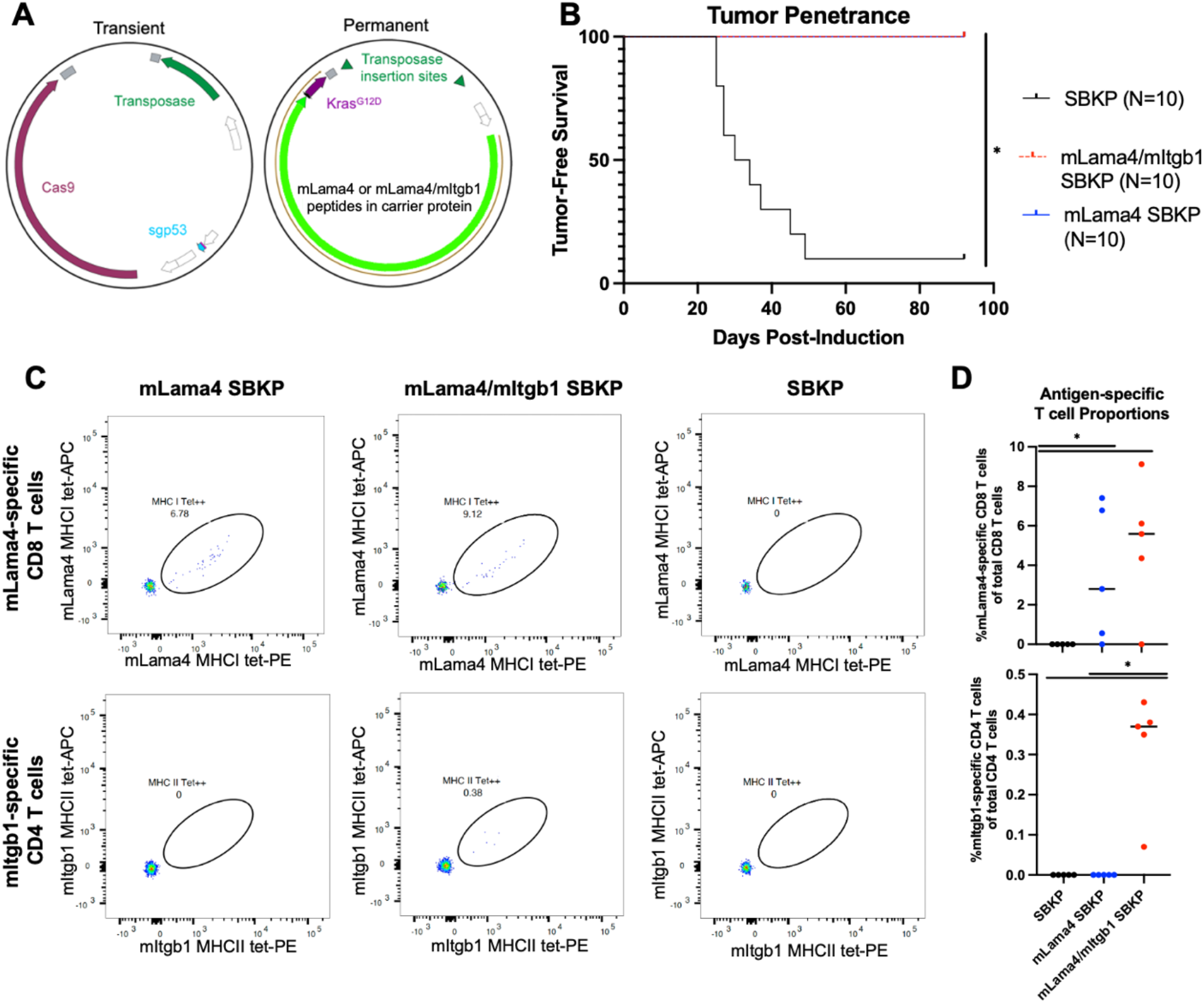
IVE of SBKP plasmids containing sarcoma neoantigens fails to yield tumors and results in a tumor-specific immune response at 10 days post-induction. **A)** Illustrations of the transient and persistent plasmids injected into the gastrocnemius muscle prior to electroporation. White segments indicate promoter regions and grey segments indicate poly-A tails. The transient plasmid is the same for all models. The persistent plasmid contains either Kras^G12D^ alone (SBKP), mLama4-2A-Kras^G12D^ (mLama4 SBKP), or a carrier protein containing the core antigenic peptides of mLama4/mItgb1-2A-Kras^G12D^ (mLama4/mItgb1 SBKP). **B)** Kaplan-Meier curve demonstrating tumor penetrance over time for mice induced with SBKP, mLama4 SBKP, or mLama4/mItgb1 SBKP plasmids. The survival curves are superimposed at 100% survival for mLama4 SBKP and mLama4/mItgb1. Statistical significance was determined with a Log-rank test (*, p<0.05). **C)** Representative flow plots depicting mLama4-specific CD8 T cells (top row) or mItgb1-specific CD4 T cells (bottom row) in PBMCs collected from mice 10 days after IVE. All samples were stained with MHC tetramers (tet) conjugated to 2 different fluorophores (PE and APC). **D)** Percent of antigen-specific T cells of the parent T cell population (CD4 or CD8) in PBMCs collected from mice 10 days after IVE. Statistical significance was determined with a non-parametric ANOVA with FDR correction for multiple comparisons (*, p<0.05).

While the expression status of these neoantigens in the original primary tumor is unknown (i.e., it is possible that these mutations were acquired *in vitro* after the cell line was established), we hypothesized that utilizing these neoantigens in an autochthonous sarcoma model was less likely to elicit a robust immune response compared to sarcomas expressing highly immunogenic model antigens such as ovalbumin or LCMV glycoproteins. While the transient plasmid remained unchanged from the SBKP model, we modified the integrating plasmid to include the known sarcoma neoantigens *mLama4* and *mItgb1* (**Figure 2A**). Specifically, we developed two integrating plasmid variants. The first integrating plasmid (mLama4 SBKP) contained the entire mLama4 protein coding sequence followed by a 2A sequence leading into *Kras*^*G12D*^. The second integrating plasmid (mLama4/mItgb1 SBKP) contained a GFP carrier protein with the verified core binding peptides of both mLama4 and mItgb1 followed by a 2A sequence leading into *Kras*^*G12D*^ (**Figure 2A**). Neoantigens were intentionally placed upstream of *Kras*^*G12D*^ and on a single transcript to minimize the chances of selective silencing of the neoantigen in tumor cells, as loss of neoantigen expression would also inhibit expression of the driver oncogene *Kras*^*G12D*^.

When electroporated along with the transient plasmid containing Cas9, sgRNA to *Trp53*, and sleeping beauty transposase, both of these integrating plasmids transformed MEFs as measured by similar growth in soft agar as compared to the SBKP-integrating plasmid, which lacked expression of neoantigens on the integrating plasmid (**Supplemental Figure 1**). Yet, after IVE into the gastrocnemius of immunocompetent C57BL/6J wildtype mice, tumors failed to form after ∼3 months (**Figure 2B**). Using MHC tetramer flow cytometry of PBMCs, we identified mLama4-specific CD8 T cells as early as 10 days after IVE in both the mLama4 SBKP and mLama4/mItgb1 SBKP models, but not in the SBKP control model (**Figure 2C; Supplemental Figure 2**). A detectable mItgb1-specific CD4 T cell response was also observed specifically in mice electroporated with the mLama4/mItgb1 SBKP integrating plasmid, but in none of the mice electroporated with an integrating plasmid lacking mItgb1. In addition, PBMCs from mLama4/mItgb1 SBKP mice and mLama4 SBKP mice collected 10 days after IVE contained significantly higher relative levels of mLama4-specific CD8 T cells compared to SBKP controls (**Figure 2D**, mLama4/mItgb1 SBKP: mean % mLama4-specific CD8/total CD8 = 5.0, FDR-corrected p=0.02; mLama4 SBKP: mean % mLama4-specific CD8/total CD8 = 3.3, FDR-corrected p=0.04; SBKP: mean % mLama4-specific CD8/total CD8 = 0). Additionally, mLama4/mItgb1 SBKP mice were the only group that demonstrated detectable levels of mItgb1-specific CD4 T cells (mLama4/mItgb1 SBKP: mean % mItgb1-specific CD4/total CD4 = 0.3, FDR-corrected p=0.002). Taken together, these data are consistent with a model in which the introduced neoantigens drive an early immune-mediated clearance of tumor-initiating cells that prevents gross tumor formation.

Although the soft agar assay confirmed the ability of the neoantigen-containing SBKP plasmids to transform cells *in vitro* (**Supplemental Figure 1**), to test the ability of the neoantigen-containing SBKP plasmids to initiate sarcomagenesis *in vivo*, we tested whether the plasmids could induce sarcomas *in vivo* in the absence of CD4 and CD8 T cells. We systemically depleted CD4 and CD8 T cells 3 days prior to IVE of the mLama4/mItgb1 SBKP, mLama4 SBKP, and SBKP plasmid sets. Under these conditions, mice from all groups developed gross tumors at ∼30 days with statistically similar incidence of tumor-free survival (**Figure 3A**, p=0.1). To confirm that mLama4 was still expressed in the resulting tumors of the mLama4/mItgb1 SBKP group, sarcoma cell lines were generated from primary tumors and transplanted into naive C57BL/6J mice. PBMCs of transplanted mice were collected 7 days after transplantation and subjected to MHCI tetramer flow cytometry to identify CD8 T cells specific for mLama4 (**Figure 3B**). Transplanted mice demonstrated a mean proportion of mLama4-specific CD8 T cells of total T cells of 1.4% (**Figure 3C**, p=0.003). The development of mLama4-specific CD8 T cells confirmed that the neoantigen was still present in the primary tumor cells. Taken together, these results establish that the neoantigen-containing SBKP plasmids are competent to initiate sarcomagenesis *in vivo* in the absence of CD4 and CD8 T cells. Therefore, these results suggest that the lack of sarcoma development following IVE of the neoantigen-containing SBKP plasmids in immunocompetent C57BL/6J wildtype mice is a consequence of lymphocyte-mediated immune clearance.

**Figure 3.**
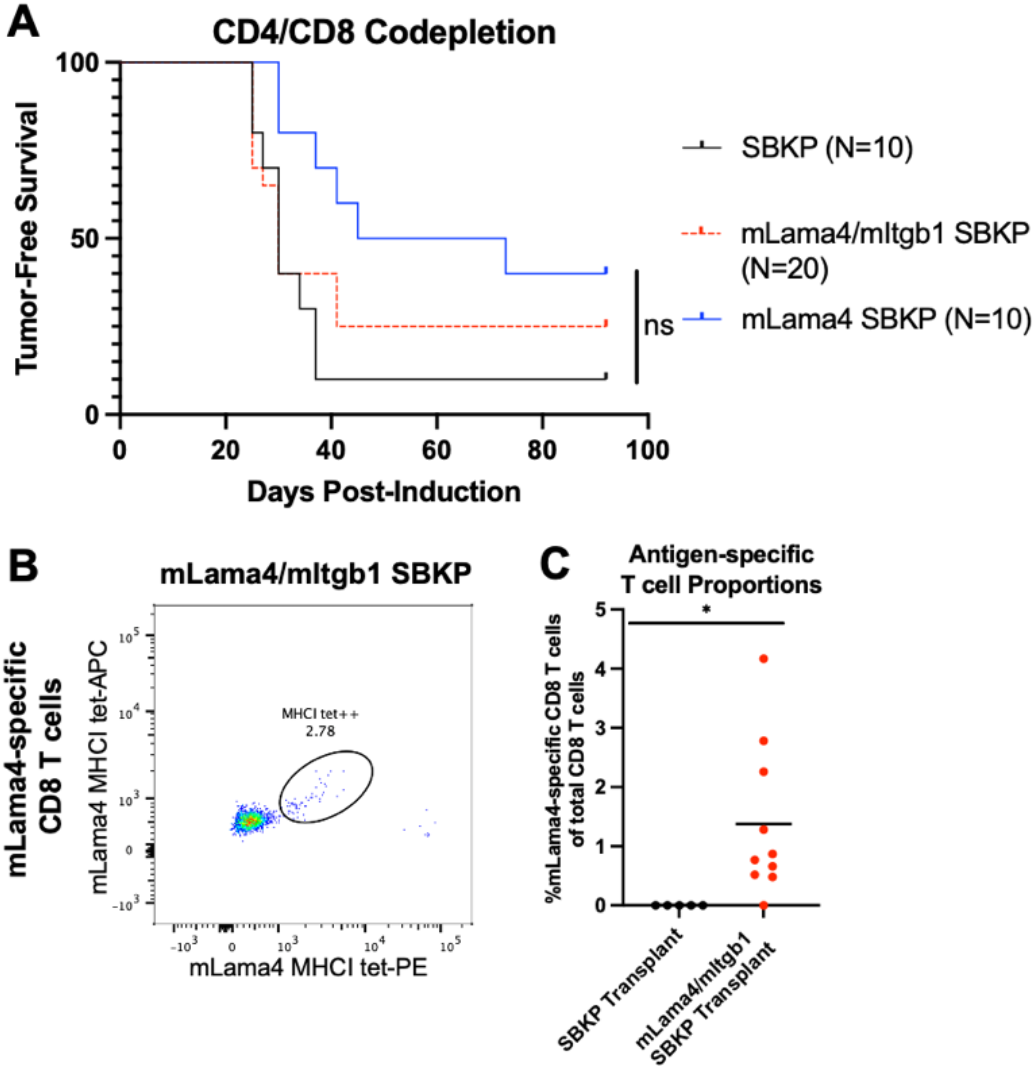
Depletion of CD8 and CD4 T cells prior to IVE permits sarcoma neoantigen-expressing tumor formation that persists throughout tumor development. **A)** Kaplan-Meier curve of tumor-free survival after mice underwent IVE to the gastrocnemius muscle with the SBKP, mLama4/mItgb1 SBKP, or mLama4 SBKP plasmid systems. Mice were injected once with anti-CD4 and anti-CD8 antibodies to deplete T cells 3 days prior to IVE. Statistical significance was determined with a Log-rank test. After collection of mLama4/mItgb1 autochthonous tumors, cells were transplanted into syngeneic recipient C57BL/6J mice (N=10). **B)** Representative flow plot depicting mLama4-specific CD8 T cells in PBMCs collected from a recipient mouse 7 days after transplantation of mLama4/mItgb1 SBKP tumor cells. All samples were stained with MHC tetramers conjugated to 2 different fluorophores (PE and APC). **C)** The percent of mLama4-specific T cells of total CD8 T cells in the mice 7 days after transplant with mLama4/mItgb1 SBKP tumor cells (N=10) compared to SBKP tumor cells (N=5). Statistical significance was determined with an unpaired non-parametric T test.

### CD4 and CD8 T cells contribute to early immune-mediated tumor clearance

While CD4 and CD8 T cell co-depletion permitted tumor growth in both the mLama4 SBKP and mLama4/mItgb1 SBKP models, the effect of depleting each T cell subset was unclear. In transplanted sarcoma cell lines expressing the MHCI-presented neoantigen mLama4 and/or the MHCII-presented neoantigen mItgb1, both neoantigens were required for decreased tumor penetrance and response to anti-PD-1 treatment, which indicates that in a transplanted tumor model both CD8 and CD4 T cells contribute to immune-mediated sarcoma clearance^10^. To evaluate the contribution of the CD8 and CD4 T cell compartments in an autochthonous sarcoma setting where the tumor co-evolves with the immune system, we electroporated mice with plasmids after depleting CD4 T cells alone, CD8 T cells alone, both T cell subsets, or neither T cell subset (**Figure 4**). As predicted, in the SBKP model that does not contain any known tumor neoantigens, depletion of the different T cell subsets had little effect on tumor penetrance (**Figure 4A**). In mice induced with the mLama4/mItgb1 SBKP system, co-depletion of CD4 and CD8 T cells was necessary for high tumor penetrance (tumor penetrance=75%; **Figure 4B**). Only 10% of these animals developed a tumor after depletion of the CD8 T cell subset alone, and no animals developed tumors after depletion of the CD4 T cell subset alone. These results indicate that CD8 and CD4 T cells each contribute to immune clearance in an autochthonous model tumor system. Interestingly, while the mLama4 SBKP model demonstrated the highest tumor penetrance after CD8 and CD4 T cells co-depletion (tumor penetrance=60%), tumors did form after CD4 T cell depletion alone in 20% of the animals (**Figure 4C**). Given that the mLama4 SBKP plasmid does not contain a known MHCII-presented neoantigen, the ability of CD4 T cell depletion to bypass immune surveillance to permit sarcoma development was unexpected. It is conceivable that an unidentified MHCII-presenting neoantigen is present within the complete coding sequence of mLama4 that contributes to a helper T cell response, which enhances CD8 T cell killing. Alternatively, CD4 T cells could be inhibiting tumor growth through alternative mechanisms independent of CD8 T cells, such as NK cell recruitment.

**Figure 4.**
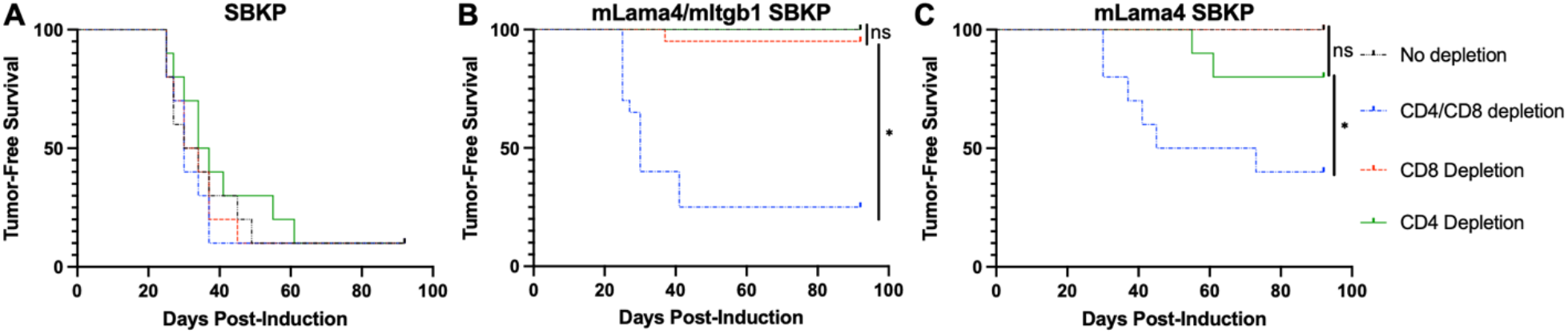
CD4 and CD8 T cells are both independently capable of clearing early neoantigen-containing autochthonous sarcomas in mice following IVE. **A)** SBKP (N=10 per depletion condition), **B)** mLama4/mItgb1 SBKP (N=20 per depletion condition), and **C)** mLama4 SBKP tumors (N=10 per depletion condition) were induced with IVE 3 days after systemic depletion with isotype control antibody, anti-CD4, anti-CD8, or both. Kaplan-Meier curves demonstrate tumor penetrance over time. Statistical significance was determined with a Log-rank test (ns, not significant).

## Discussion

In this study, we developed a novel model of autochthonous sarcoma in mice that presents the known sarcoma neoantigens, mLama4 and mItgb1^10,13^. This model can be used to study mechanisms of immunosurveillance during the early steps of tumor formation. Here, we used this autochthonous model to test the requirement of CD8 and CD4 T cells in immunosurveillance. Several studies in mice and humans have demonstrated that tumor antigen-specific immune cell populations can differ dramatically from bulk tumor-infiltrating immune cells, both at baseline and in response to therapy^10,21^. In these cases, significant phenotypic responses of tumor-specific T cells are diluted by the lack of responses in the tumor non-specific T cell population. We selected the sarcoma-derived neoantigens mLama4 and mItgb1 for this model in an effort to moderate the immunogenicity compared to frequently studied model antigens such as LCMV glycoproteins or ovalbumin, which are more highly immunogenic than typical tumor neoantigens in humans. Ideally, an autochthonous sarcoma model with known antigen expression would develop without requiring depletion of T cells because immune cell depletion disrupts the coevolution between the tumor and immune system, which is one of the key benefits of autochthonous tumor models over transplant models. However, this mLama4- and mItgb1-expressing autochthonous tumor model fails to result in sarcoma development in mice with a fully intact immune system due to both CD4 and CD8 T cell responses.

To address the limitation of this model, in future studies modifications to the integrating plasmid could be made to try to preserve tumor-immune system coevolution, while potentially permitting tumor penetrance. For example, the expression of oncogenic Kras^G12D^ could be decoupled from the neoantigen on the integrating plasmid so that a doxycycline-induced regulatory element could be added to temporally restrict neoantigen expression until tumors have started to develop. This approach would not only provide additional time that may allow the tumor to outcompete the immune system before activating neoantigen expression, but would also provide an opportunity to tune the amount of neoantigen expression according to the concentration of doxycycline. Decreasing expression of the neoantigens could allow tumor cells to escape immunosurveillance so that sarcomas develop without T cell depletion. An alternative approach to potentially decrease the expression of the neoantigens would be to insert the neoantigen cassette into the genome after the endogenous promoter of oncogenic *Kras*^*G12D*^. A limitation inherent to IVE of multiple plasmids is the difficulty controlling the multiplicity of integration; different cells will likely receive disparate copy numbers of each plasmid. In our model, this may result in high neoantigen expression levels that enhance an early anti-tumor immune response leading to tumor cell clearance.

While the anti-tumor function of CD8 T cells is relatively clear, the potential for CD4 T cells to act as anti-tumor inflammatory helper cells or pro-tumor immunosuppressive regulatory cells makes the relationship of CD4 T cells to tumor development, growth, and therapeutic response less straightforward. Tregs developing from naïve CD4 T cells in the tumor draining lymph nodes have been reported to promote immune tolerance^6^. Yet, a growing body of evidence has also emerged to uncover an array of CD4 T cell anti-tumor mechanisms including canonical CD8 T cell help^9^, direct killing of MHCII-expressing tumor cells^22,23^, and induction of cancer cell senescence^24^. In transplant tumor models, an important anti-tumor role of CD4 T cells has been reported. Sarcoma cells expressing mLama4 or mItgb1 alone readily grew into anti-PD-1-resistant tumors after transplantation into mice. Yet, coexpression of both the MHCI- and MHCII-presenting mLama4 and mItgb1 neoantigens led to spontaneous tumor regression in most transplanted mice and a clear anti-PD-1 response in the remaining tumors that did continue to grow^10^. These results suggest that recruitment of either CD8 T cells or CD4 T cells alone is insufficient to promote an effective antitumor response and that recruitment of both T cell subsets greatly enhances the anti-tumor immune response. We sought to interrogate the importance of the tumor-specific CD8 and CD4 T cell subsets in an autochthonous tumor model using the same neoantigens expressed after IVE-induced autochthonous sarcoma development. CD8 and CD4 T cell co-depletion was sufficient and largely necessary for tumor growth, as depletion of either T cell subset alone (i.e. CD8 or CD4 T cells) led to tumor formation in only a few animals. This suggests that either CD8 or CD4 T cell responses were sufficient for immune-mediated tumor clearance in the autochthonous setting. While CD8 T cells are capable of transitioning from naïve into activated effectors without CD4 help, the mechanisms of CD4 T cell-mediated tumor clearance in the absence of CD8 T cells in this model remains unclear. It is possible that CD4 T cells stimulated an anti-tumor macrophage response, tumor cell senescence, and/or promoted tumor vasculature degradation. Interestingly, even in the mLama4 SBKP model, which lacks a known MHCII neoantigen, tumors failed to grow after CD8 T cell depletion. It is possible that an undefined MHCII neoantigen is present in this model, which leads to a CD4 T cell-mediated anti-tumor effect. Perhaps tissue injury induced be IVE could promote the mutations in tumor initiating cells that lead to the expression of additional tumor neoantigens.

In conclusion, we developed a novel neoantigen-bearing autochthonous sarcoma model to study tumor-specific immune cell responses. Due to strong immunosurveillance, T cell depletion was required for high tumor penetrance. While studies using transplant sarcomas have reported that both CD4 and CD8 T cell responses are necessary for tumor clearance and therapeutic responses, our data in the autochthonous model suggests that CD4 and CD8 T cells are each capable of tumor clearance independently of the other T cell subset. These findings support further research into anti-tumor mechanisms of CD4 T cells as they could prove therapeutically valuable.

Furthermore, these data provide another example where the immune response to autochthonous tumor models does not recapitulate the immune response of transplant tumor models. We anticipate that the models developed in this study will be valuable tools to further explore tumor-specific immune cell mechanisms of tumor clearance.

## Acknowledgements

We thank the NIH Tetramer Core Facility for providing the MHC tetramers used in this study. This work was supported by 2R35 CA197616 (DGK) from the National Cancer Institute, The Leon Levine Foundation (DGK), and the Merck Investigator Studies Program (DGK).

## Supplemental Figures/ Tables

**Supplemental Table 1**

**Supplemental Figure 1**

**Supplemental Figure 2**

