## Supplemental Figures/ Tables for "Both CD8 and CD4 T cells contribute to immunosurveillance preventing the development of neoantigen-expressing autochthonous sarcomas"

### Slide 1
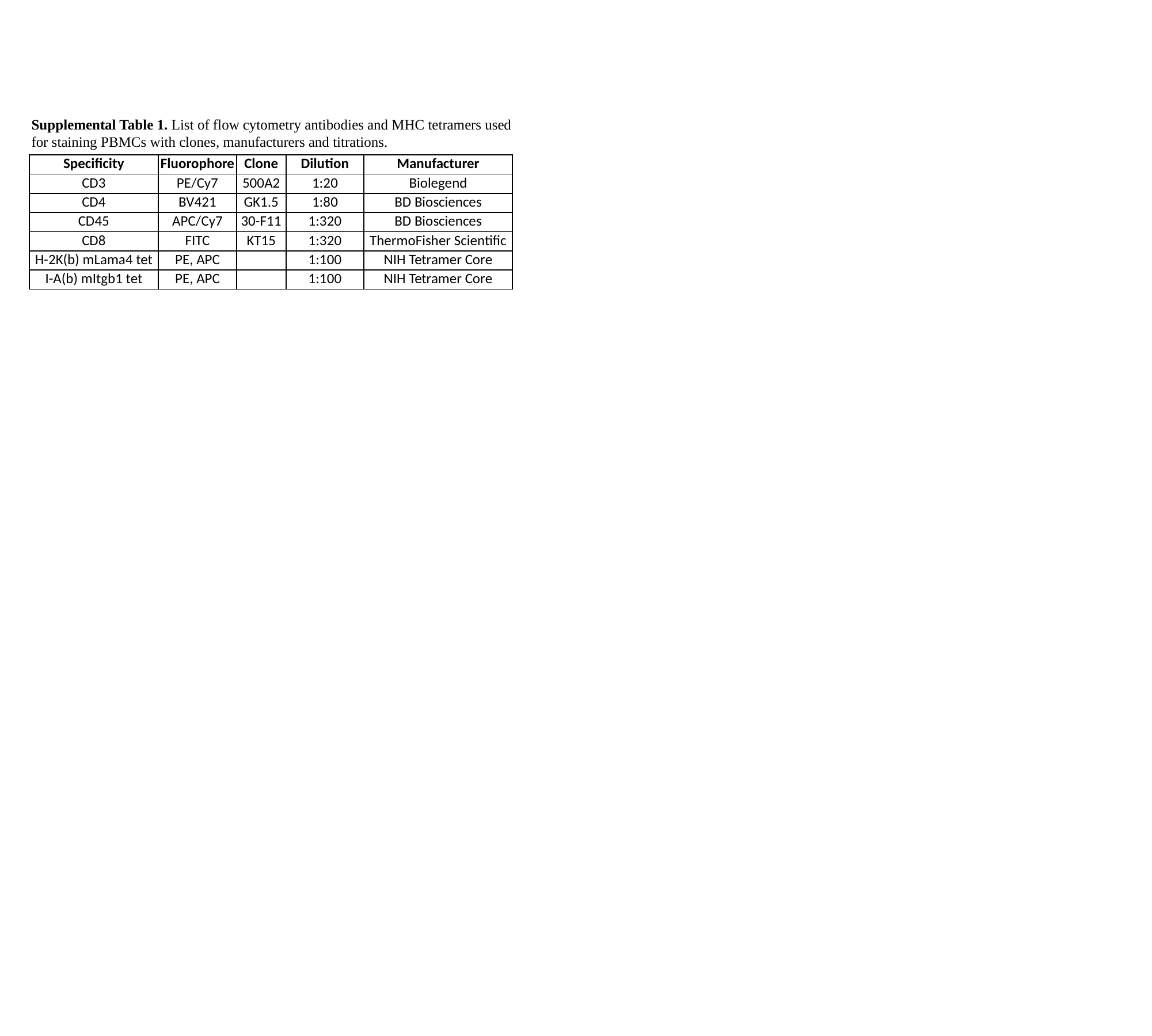

Supplemental Table 1. List of flow cytometry antibodies and MHC tetramers used for staining PBMCs with clones, manufacturers and titrations.
| Specificity | Fluorophore | Clone | Dilution | Manufacturer |
| --- | --- | --- | --- | --- |
| CD3 | PE/Cy7 | 500A2 | 1:20 | Biolegend |
| CD4 | BV421 | GK1.5 | 1:80 | BD Biosciences |
| CD45 | APC/Cy7 | 30-F11 | 1:320 | BD Biosciences |
| CD8 | FITC | KT15 | 1:320 | ThermoFisher Scientific |
| H-2K(b) mLama4 tet | PE, APC | | 1:100 | NIH Tetramer Core |
| I-A(b) mItgb1 tet | PE, APC | | 1:100 | NIH Tetramer Core |

### Slide 2
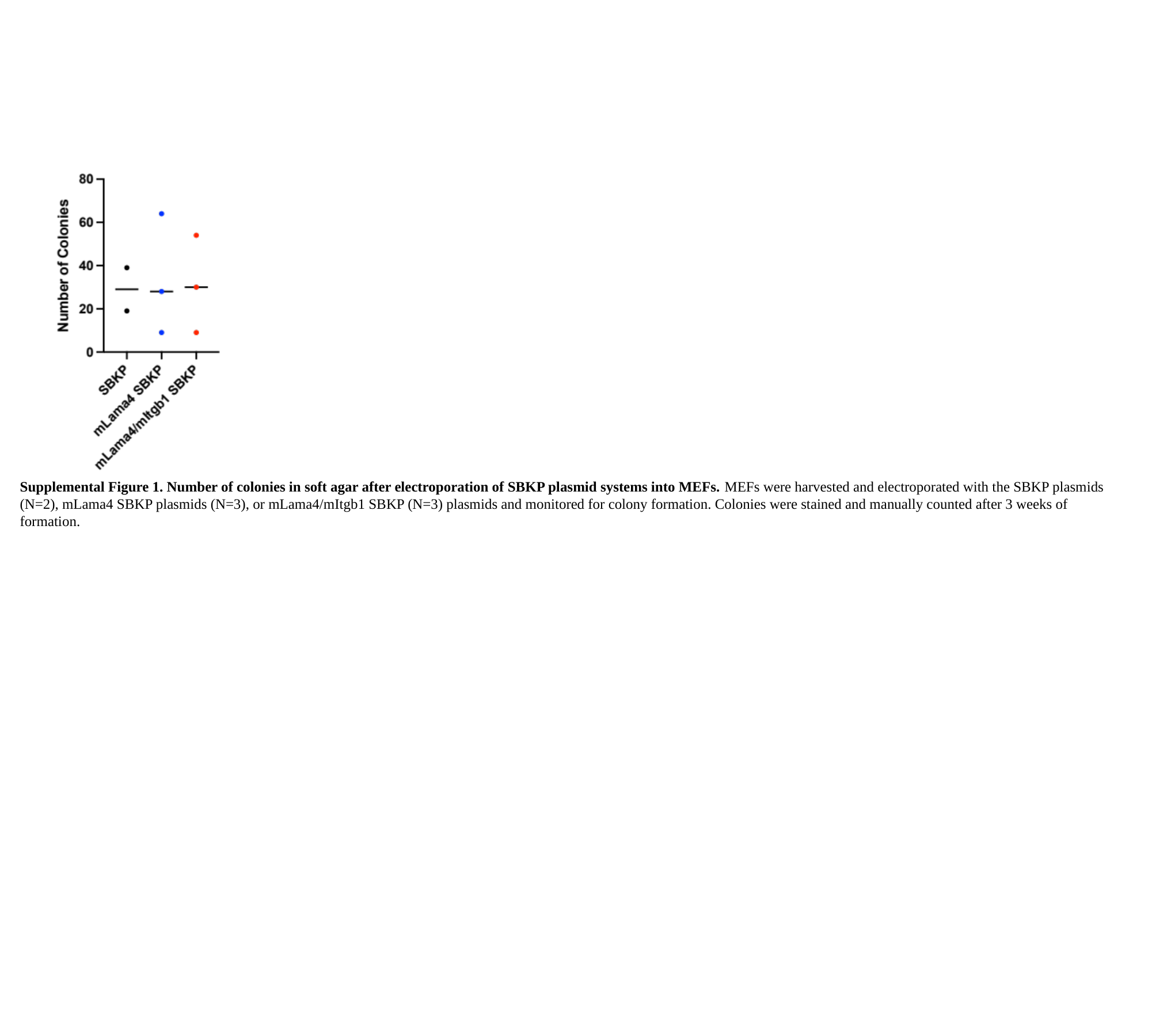

Supplemental Figure 1. Number of colonies in soft agar after electroporation of SBKP plasmid systems into MEFs. MEFs were harvested and electroporated with the SBKP plasmids (N=2), mLama4 SBKP plasmids (N=3), or mLama4/mItgb1 SBKP (N=3) plasmids and monitored for colony formation. Colonies were stained and manually counted after 3 weeks of formation.

### Slide 3
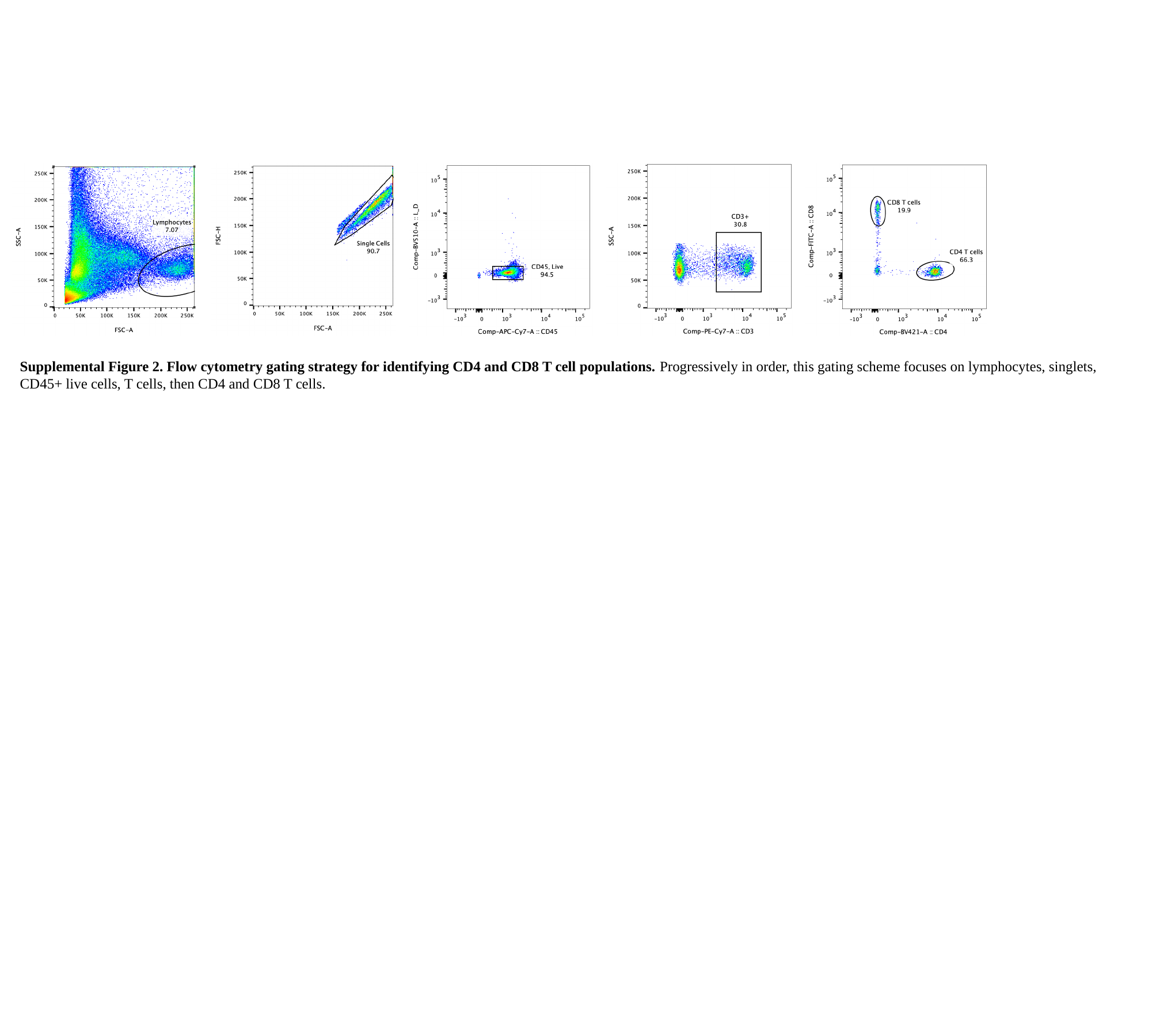

Supplemental Figure 2. Flow cytometry gating strategy for identifying CD4 and CD8 T cell populations. Progressively in order, this gating scheme focuses on lymphocytes, singlets, CD45+ live cells, T cells, then CD4 and CD8 T cells.
